# Extended practice improves the accuracy and efficiency of goal-directed reaching guided by supplemental kinesthetic vibrotactile feedback

**DOI:** 10.1101/2022.05.18.491184

**Authors:** Valay A Shah, Ashiya Thomas, Leigh A Mrotek, Maura Casadio, Robert A Scheidt

**Affiliations:** Dept. Biomedical Engineering, Marquette University and Medical College of Wisconsin, Milwaukee, WI, USA, 53233; Dept. Aging and Geriatric Research, University of Florida, Gainesville, FL, USA, 32611; DIBRIS, University of Genova, Genova, Italy, 16145

**Keywords:** Practice, sensory augmentation, dual-tasking, sensorimotor learning

## Abstract

Prior studies have shown that providing task-specific vibrotactile feedback (VTF) during reaching and stabilizing with the arm can immediately improve the accuracy and efficiency. However, such studies typically evaluate performance after less than 1 hour of practice using VTF. Here we tested the effects of extended practice using supplemental kinesthetic VTF on goal-directed reaching with the arm. Healthy young adults performed a primary reaching task and a secondary choice reaction task individually and as a dual-task. The reaching task was performed under three feedback conditions: visual feedback, proprioceptive feedback, and with supplemental kinesthetic VTF applied to the non-moving arm. We compared performances before, during, and after approximately 10 hours of practice on the VTF-guided reaching task, distributed across 20 practice sessions. Upon initial exposure to VTF-guided reaching, participants were immediately able to use the VTF to improve reaching accuracy. Performance improvements were retained from one practice session to the next. After 10 hours of practice, the accuracy and temporal efficiency of VTF-guided reaching were equivalent to or better than reaching performed without vision or VTF. However, hand paths during VTF-guided reaching exhibited a persistent strategy whereby movements were decomposed into discrete sub-movements along the cardinal axes of the VTF interface. Dual-tasking capability also improved, such that the primary and secondary tasks we performed more concurrently after extended practice. Our results demonstrate that extended practice on VTF-guided reaching can yield performance improvements that accrue in a manner increasingly resistant to dual-task interference.

## Introduction

When a person moves through a complex environment, they typically use information from multiple senses - including vision and proprioception - to guide their ongoing actions (Sober and Sabes 2003; Scott 2004). Neural injuries such as those caused by stroke can disrupt sensorimotor pathways used for controlling movements vital to an independent lifestyle (Carey 1995; Hermsdörfer et al. 2003; Zackowski et al. 2004; Scheidt and Stoeckmann 2007; Blennerhassett et al. 2007; Tyson et al. 2008; Dukelow et al. 2010; Carey and Matyas 2011; Carlsson et al. 2018). Whereas motor impairments are common and conspicuous, impaired somatosensation of limb position and movement [i.e., impaired *kinesthesia* (Gharbawie et al. 2005; Sullivan et al. 2011; Proske and Gandevia 2012)] also frequently occurs. Recent experimental studies find that more than 50% of survivors of stroke experience kinesthetic deficits in their contralesional limbs (Dukelow et al. 2010; Semrau et al. 2013). Kinesthesia is critical for the planning and control of movement and kinesthesia plays an important role in the recovery of arm function after neuromotor injury (Wade et al. 1983; Sainburg et al. 1993; Turville et al. 2017). To compensate for somatosensory deficits, people tend to rely heavily on visual feedback to control ongoing movements (Bonan et al. 2004). But visual feedback takes longer to process than somatosensation, delaying detection and correction of performance errors, thereby leading to slow and jerky movements (Sarlegna et al. 2006; Cameron et al. 2014). Many stroke survivors give up using their contralesional limb due to their impaired somatosensation despite retaining some ability to move their limb, and even though this reduces quality of life. To what extent might it be possible to circumvent injured sensorimotor control pathways and re-establish closed-loop control of the more-involved arm and hand?

This paper takes initial steps toward addressing that question by determining the extent to which extended practice with a novel wearable sensory feedback device can promote improved accuracy and efficiency of goal-directed arm movements in a small cohort of neurologically intact individuals. The approach uses movement sensors attached to one arm and a non-invasive sensory interface attached elsewhere on the body to deliver supplemental feedback of hand location, which can be used to improve the accuracy and efficiency of unseen reaching movements. For this approach to work, the system’s user must learn a novel mapping between the state of the moving limb and changes in the synthesized feedback delivered by the sensory interface, and then use that map to guide subsequent actions [cf. König et al. (2016)]. Preliminary studies have demonstrated that within minutes of initial exposure to a novel sensory interface, neurologically intact individuals can rapidly learn to use supplemental feedback of limb configuration and movement to improve the accuracy of reaching actions performed without visual guidance and some survivors of stroke (Tzorakoleftherakis et al. 2015, 2016; Krueger et al. 2017; Risi et al. 2019; Ballardini et al. 2021).

The idea of providing supplemental feedback to mitigate sensory loss (sensory substitution) or impairment (sensory augmentation) has been explored for decades (Gault 1926a; Bach-Y-Rita et al. 1969; White et al. 1970). Vibrotactile sensory interfaces show particular promise for enhancing postural stabilization in vestibular patients and for providing information about grasp force and hand aperture to users of myoelectric forearm prostheses (Sienko et al. 2008; Lee et al. 2011, 2012; Witteveen et al. 2015). While others have also proposed vibrotactile sensory interfaces to train body motions by providing feedback of spatial orientation or by stimulating the moving limb in response to limb configuration, few have attempted to mitigate the effects of somatosensory impairment or loss on the control of contralesional arm and hand movements after neuromotor injury errors (Lieberman and Breazeal 2007; Kapur et al. 2009; Weber et al. 2011; Bark et al. 2015; König et al. 2016). Those that have attempted this were largely technology demonstrations or feasibility case studies (De Santis et al. 2014; Afzal et al. 2015; Hussain et al. 2015; Elangovan et al. 2019; Ballardini et al. 2021). None have examined the extent to which extended practice with such technology can facilitate learning of the novel sensorimotor relationships needed to establish novel closed-loop feedback control of a moving limb [but see König et al. (2016) for a training study of sensory augmentation in the context of spatial navigation].

Here, we tested the extent to which extended practice can promote skilled use of supplemental vibrotactile kinesthetic feedback to control arm movements performed without ongoing visual feedback. According to classical descriptions, a skill becomes increasingly resistant to dual-task interference as it is learned [cf., Fitts and Posner (1967)]. We implemented a dual-task experimental paradigm designed to determine the extent to which practice using a novel wearable sensory prosthesis can enhance performance in a manner resistant to dual-task interference. We asked healthy, young adults to practice using real-time vibrotactile feedback (VTF) of hand location to guide reaching in movements in a 2D workspace. Participants trained for 10 hours each over the course of four weeks. We analyzed performance in the primary reaching task before, during, and after practice. We employed a secondary choice reaction task to probe constraints on attentional resources during reaching and the extent to which the primary task is automatized as skill is acquired and its demands on attentional resources are reduced. We expect our results will show improved accuracy and efficiency of goal-directed reaching after extended practice using limb-state VTF.

## Methods

Fifteen neurologically intact participants (age: 23 to 28 years; eight female) participated in this study after providing written informed consent approved by Institutional Review Boards serving Marquette University and the University of Genoa in accordance with the Declaration of Helsinki (Protocol Number: HR-3044). All participants were naïve to vibrotactile feedback (VTF) and to the study objectives, and none had known cognitive deficits or tactile deficits of the arm. Eleven participants self-identified as right-handed while four identified as left-handed. Each participant completed a total of 20 experimental sessions lasting less than 60 minutes each. Sessions were spaced at least six hours apart, with no more than two sessions per day and no more than five sessions per week. During the sessions, participants completed a primary goal-directed reaching task and a secondary choice reaction time task under various feedback conditions, as described below.

### General Experimental Setup

Each participant sat in an adjustable height chair and used their dominant hand to grasp the handle of a passive two-degree-of-freedom planar manipulandum placed directly in front of them such that the device’s entire workspace was comfortably within reach (Fig 1A). The passive manipulandum comprised a pantographic linkage with integrated potentiometers that provide signals uniquely determined by hand position in the horizontal plane [see Ballardini et al. (2018) for full design specifications]. Hand position data were collected at a rate of 50 samples per second using a custom script within the MATLAB computing environment (version R2017a; the MathWorks Inc., Natick MA). A visual display screen (LG Inc, Model: 23EA63V-P, 60 Hz) was placed vertically in front of the participant just beyond the manipulandum (Fig 1A). Visual stimuli were created and displayed in real-time using PsychToolbox-3 for MATLAB (Brainard 1997). These included a set of 25 reaching targets (0.5 cm diameter circles) arranged in a 5×5 grid on the vertical screen. Inter-target distance was 2 cm and the target grid was centered on the vertical screen (Fig 1B). Hand position on the manipulandum was mapped onto the position of a cursor (0.5 cm diameter disk) on the vertical display screen in a 1:1 ratio, such that 1 cm movement on the manipulandum resulted in a 1 cm movement of the cursor on the screen. An opaque drape blocked direct view of the dominant arm and manipulandum. The participant wore noise-cancelling headphones that played white noise to minimize extraneous auditory cues.

**Figure 1:**
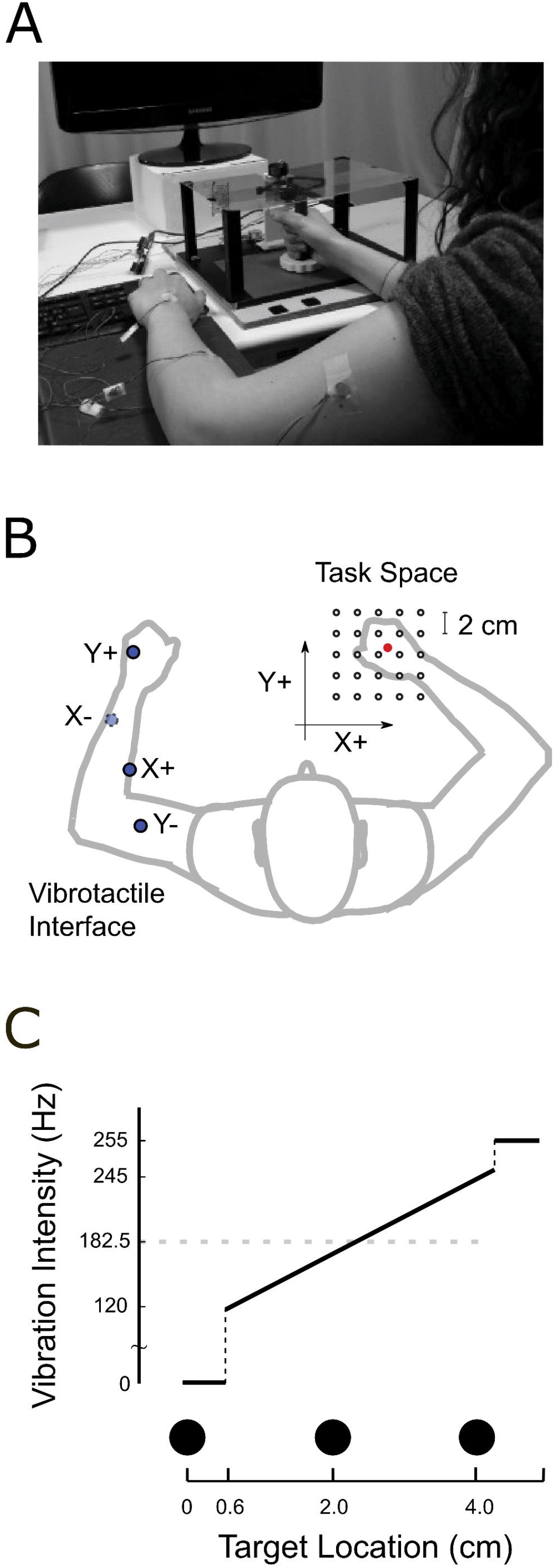
Experimental setup. **Panel A)** The participant grasped the handle of a horizontal planar manipulandum with the dominant hand. A vertical monitor was placed behind the manipulandum and displayed the visual reaching targets. The non-moving arm was rested on a rigid support with the vibrotactile interface attached via tape. **Panel B)** Vibrotactile interface made of four ERM motors (nominal locations indicated by *blue* markers) attached to the non-moving arm while the moving arm works in a 5×5 grid of reaching targets (*red* symbols). **Panel C)** Illustration of the piecewise mapping between hand displacement and vibration motor activation that defined the VTF interface. Black circles represent the location of three targets arranged horizontally. The circle on the left represents the center target. VTF is not engaged until the hand is 0.6 cm from the middle of the center target. Between 0.6 cm and 4.0 cm vibration intensity increases at a linear rate. Vibration saturates at distances greater than 4.0 cm from the center of the workspace. This pattern was followed for each of the four vibration motors.

The participant’s stationary, non-dominant arm rested on a rigid support structure with the elbow at 90° of flexion and the shoulder at 45° abduction. We placed the numeric keypad of a standard computer keyboard comfortably under the fingers of this hand. We covered three keycaps of the numeric keypad with colored tape (red, blue, green) such that the participant could use their index, middle, and ring fingers to perform a secondary, choice reaction time task based on colored visual stimuli, as described below.

A vibrotactile sensory interface was attached to four locations on the non-moving (non-dominant) arm to provide supplemental kinesthetic feedback of the moving hand’s location in the workspace (Fig 1A). The interface comprises four eccentric rotating mass vibration motors (ERM; Precision Microdrives Inc, Model: 310-117). The vibration motors are powered and controlled using custom drive circuitry interfaced to the experimental computer running MATLAB. The vibration motors provided feedback of the dominant arm’s hand position relative to the center of the workspace (Krueger et al. 2017; Risi et al. 2019). Vibration motors providing information about position in the left/right dimension were placed on the dermatome C7 and T1 on the forearm (Fig 1B: X+ and X-). Vibration motors providing information about the anterior/posterior dimension were placed on the dorsum of the hand and dermatome C5 on the skin above the lateral biceps (Fig 1B: Y+ and Y-).

The vibrotactile interface provides continuous supplemental kinesthetic feedback about hand position through changes in vibration intensity. The position of the manipulandum’s handle (and thus the position of the hand in the 2D workspace) was converted to vibration intensity in a piecewise linear manner. Vibration was turned off when the cursor was within the boundary at the center target of the workspace, vibration intensity increased discontinuously to 120 Hz when the hand moved outside of the central target, and vibration intensity increased linearly (30 Hz /cm) until it reached a maximum intensity of 255 Hz at distances greater than 4.5 cm from the center of the display (Fig 1C). Note that a 2 cm inter-target distance yields an inter-target change in vibration intensity of 60 Hz, which was greater than the average just noticeable difference in vibration intensity (JND = 37 Hz) identified for dermatomes of the arm and hand in healthy adults (Shah et al. 2019a). The 2.0 cm inter-target distance also is below the threshold of limb position acuity (2.5 ± 0.20 cm) derived using joint angular uncertainty values reported by Fuentes and Bastian (2009). We intended these experiment design choices to increase the likelihood that participants would rely on the supplemental VTF rather than intrinsic proprioception to accurately perform the primary reaching task described in the following paragraphs.

### Tasks

Participants performed two tasks, a primary reaching task and a secondary choice reaction task in blocks of 25 trials each. Both tasks were performed separately (single-task conditions) as well as concurrently in a dual-task context (Table 1). Participants performed different combinations of reaching task blocks, choice reaction task blocks, and dual-task blocks depending on the session number (Table 1). Participants received ∼ 30 minutes of practice on VTF-guided reaching during all 20 sessions (i.e., during trial blocks 6-10) post-practice assessment of dual-task performance was measured during the 1^st^, 10^th^, and 20^th^ sessions (block 11).

**Table 1:** Block schedule and feedback conditions within each trial block of each session.

| Block | Trials | Task | Feedback | Session | Type |
| --- | --- | --- | --- | --- | --- |
| 1 | 25 | RCH | V+T- | 1 | Baseline |
| 2 | 25 | RCH | V-T- | 1 | Baseline |
| 3 | $\geq 25$ | CRT | N/A | 1,10,20 | Familiarization |
| 4 | $\geq 25$ | RCH | V+T+ | 1-20 | Familiarization |
| 5 | 25 | DT | V-T+ | 1 | Pre-Practice |
| 6-10 | 25/block<br>(125 total) | RCH | V-T+ | 1-20 | Practice |
| 11 | 25 | DT | V-T+ | 1,10,20 | Post-Practice |

RCH: reach task; CRT: choice reaction task; DT: dual-task including RCH and CRT performed simultaneously. V+T-: with Visual feedback but no VTF; V-T-: no Visual feedback and no VTF; V+T+: with Visual feedback and VTF; V-T+: no Visual feedback but with VTF. Note: not all trial blocks were performed in each session.

When reaching, participants were to move the manipulandum’s handle with their dominant hand as quickly and accurately as possible to capture a visual target presented on the screen as a white ring. Participants verbally indicated when they had reached the target, ending the trial. Reaching could be performed under three different feedback conditions (Table 1): with concurrent visual feedback of the cursor and no VTF (i.e., V+T-); with VTF and no cursor feedback (V-T+); or neither visual cursor feedback nor VTF (V-T-). At the end of each reach, participants were provided visual feedback of the onscreen cursor if it was not already visible (i.e., knowledge of results); they were then required to re-center the cursor on the current goal target, which became the starting location for the next reach trial. The visually-guided re-centering served to prevent proprioceptive drift from confounding participants’ perception of their hand position in the workspace [c.f., Wann and Ibrahim, (1992)]. Visual cursor feedback was then removed at the start of the trial in any block with the V-feedback.

The secondary choice reaction task required participants to use the index, middle, or ring finger of the non-dominant hand to indicate as quickly as possible via a button press on the color-coded keypad whenever the current target changed from a white to one of three colored disks (red, blue, or green). When performed as a stand-alone task, choice reaction trials were cued with the colored target appearing at a pseudorandomly-selected target location 1400 ± 700 ms after the completion of the previous trial.

Dual-tasking trials were always performed in the V-T+ feedback condition, i.e., without ongoing cursor feedback during the reach. During dual-task trials, the onscreen target started as a white ring, but changed to a colored disk (red, blue, or green) after a pseudorandom interval ranging from 400 -1100 ms. Participants were instructed to capture the target as quickly and accurately possible when the white ring target appeared, and to use their index, middle, or ring finger of the non-moving, non-dominant hand to press the color-corresponding button on a computer keypad as quickly as possible after the target changed colors. Participants were to continue reaching to the target while pressing the button if they had not yet finished reaching. As described above for the single-task reaching trials, participants verbally indicated when they had reached the target, ending the trial. This caused the cursor to re-appear and participants were to correct any target capture error by re-centering the cursor on the current target as the starting location for the next trial.

### Data Analysis

We computed three kinematic measures of movement accuracy and efficiency to assess reach performance. *Target capture error* (reaching accuracy) was defined as the absolute distance between the target location and the final position of the on-screen cursor (i.e., its position at the verbal end-of-trial indication, prior to re-centering the cursor on the target using end-of-trial visual feedback). *Target capture time* (reaching efficiency) was computed as the time difference between the start of the trial and the point in time when the participant’s hand speed fell below 10% of its maximum hand speed (just prior to participants’ verbal end-of-trial indication). We also computed *Decomposition Index* (DI), a unitless scalar efficiency metric related to how the participant chose to perform the reaching task; DI measures the extent to which hand paths move parallel to the cardinal (X, Y) axes of the vibrotactile display (which was aligned with the axes of the 2D workspace; see also the Appendix of Risi et al. (2019). DI is calculated as a weighted sum of the current change in *X* and *Y* hand position multiplied by the change in current orthogonal velocity (e.g., *X* position multiplied by *Y* velocity: 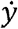). This sum was normalized to the total displacement and maximum velocity (e.g., 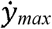) (Eq 1):

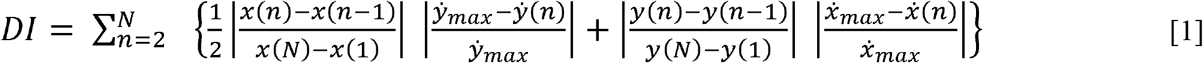

DI values increase when hand trajectories display multiple speed peaks and/or stray from a straight line connecting the start and the endpoint. Risi et al. showed that a DI value of 0.24 corresponds to straight-line trajectories with bell-shaped (minimum-jerk) velocity profiles.

To assess performance on the choice reaction task, we computed *choice reaction time* as the time difference between the moment of stimulus presentation and the button press. We computed *choice accuracy* as the percentage of correct color choices. Trials where the participant pressed the button ahead of the cue or did not press the button within the allotted trial time were recorded as errors and the reaction time from these trials was excluded from further analysis. We also analyzed the dual-task trials to determine the strategy each participant used to combine the reach and button-press tasks: we computed the percentage of trials where the participant pressed the button before the reach started, during the reach, or after the reach had ended, and we examined how the participants’ dual-task strategies changed after extended practice on VTF-guided reaching.

### Statistical Testing

We hypothesized that VTF-guided reaching would adhere to classical descriptions of sensorimotor learning whereby performance improvements accrue in a way that is increasingly resistant to dual-task interference (Fitts and Posner 1967). To test that hypothesis, we used repeated measures ANOVA and post hoc, Bonferroni-corrected paired sample *t*-tests (one-tailed) to compare task performances before (at session 1), during (at session 10), and after extended practice on VTF-guided reaching (at session 20). Dependent variables included target capture error, target capture time, DI, choice reaction time, and choice accuracy. Each dependent variable was averaged across trials within each session for each participant. Independent variables included session number and testing condition (single-task, dual-task). All analyses were performed with SPSS 27 (IBM Corp). Where appropriate, we used Greenhouse-Geisser corrections to mitigate violations of sphericity. Statistical significance was set at a family-wise error rate of α = 0.05.

## Results

This study used a dual-task experimental paradigm to evaluate practice-related improvements in the performance of planar goal-directed reaches guided by supplemental kinesthetic feedback. All participants demonstrated that they understood the primary reaching task, the secondary choice reaction task, and the dual-task instructions. They also demonstrated that they were able to use supplemental kinesthetic feedback to improve movement accuracy and efficiency of the primary reaching task. Figure 2 shows selected hand paths from movements made between the same two targets by a selected participant before/after extended practice with supplemental kinesthetic feedback. The initial hand location was the same for each movement. The straightest and most accurate reach occurred when visual cursor feedback was provided throughout the movement (Fig 2: *solid black* line). The least accurate performance occurred when participants were provided neither visual cursor feedback nor VTF (Fig 2: *dashed black* line). After brief exposure to VTF (i.e., on practice session 1), participants were immediately able to use the supplemental kinesthetic feedback to improve target capture accuracy (Fig 2: *magenta* line). After extended practice (session 20), movement accuracy was high and hand paths exhibited a strategic decomposition into discrete sub-movements along the cardinal axes of the VTF interface (Fig 2: *dashed magenta* line). As described next, performance features highlighted in Figure 2 were characteristic of the participant cohort as a whole.

**Figure 2:**
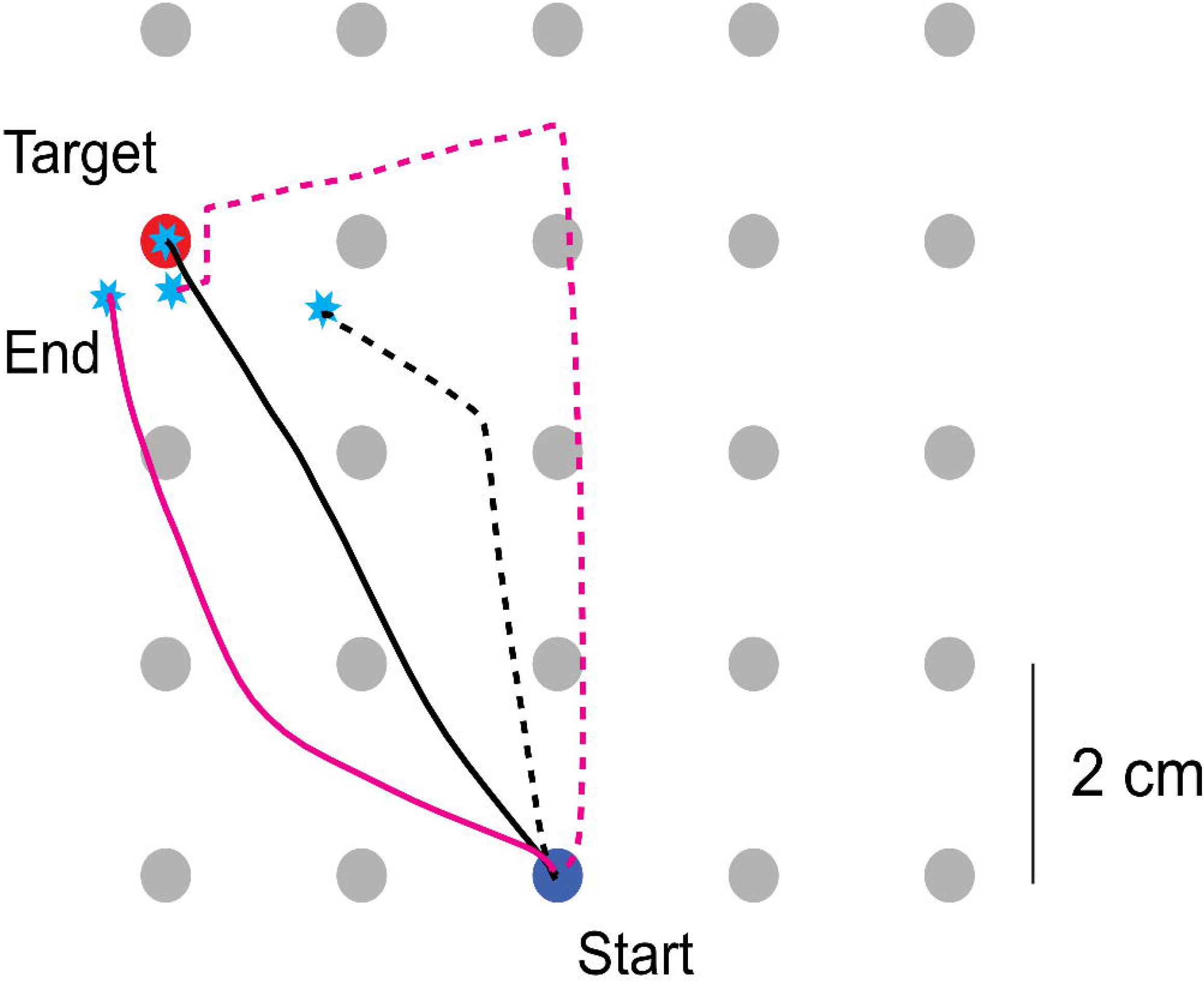
Sample reaching trajectories from one representative participant during the reaching task under various feedback conditions. Reaches were made from the *blue* start position and moved away and to the left to the *red* goal target. The *black solid* trajectory indicates visually guided reaching. The *black dashed* trajectory shows proprioceptive reaching (V-T-condition). The *magenta solid* and *magenta dashed* trajectories show pre- and post-practice VTF-guided reaching, respectively. *Cyan* stars indicate the end of each reach.

### Single-task performance in the reaching and choice reaction tasks

We analyzed three aspects of the primary reaching task under single-task conditions. First, we considered how movement accuracy changed as participants practiced reaching under the guidance of supplemental kinesthetic VTF. Figure 4 presents cohort averages from “Baseline” testing trial blocks of session 1, as well as cohort averages of mean performance in the (V-T+) “Practice” blocks performed during sessions 1 through 20. Visually guided reaching produced highly accurate movements with target capture errors averaging about 1 mm across the cohort (condition labeled “V” in Fig 3A; 0.09 ± 0.03 cm; mean ± SEM here and onward). When visual feedback was subsequently removed and participants had to rely solely on intrinsic proprioception, target capture errors increased markedly to 1.85 ± 0.37 cm (t_14_ = 18.2, p < 0.001; Fig 3A: “NV”). We used repeated measures ANOVA to examine the effect of extended practice with supplemental kinesthetic VTF on target capture error during VTF-guided reaching. This analysis revealed a main effect of practice session of movement accuracy [F_(3.16,44.2)_ = 20.7, p < 0.001]. On initial exposure to VTF, average target capture error decreased significantly from the V-T-(Fig 3A: “NV”) condition to a mean of 1.63 ± 0.30 cm in the practice blocks (t_14_ = 2.18, p = 0.023). Target capture error decreased further to 1.14 ± 0.36 cm after 10 practice sessions (t_14_ = 6.80, p < 0.001) and further still to 0.97 ± 0.25 cm after 20 practice sessions (t_14_ = 9.69, p < 0.001). The reduction in error from the 10^th^ to the 20^th^ session was also significant (t_14_ = 4.66, p < 0.001).

**Figure 3:**
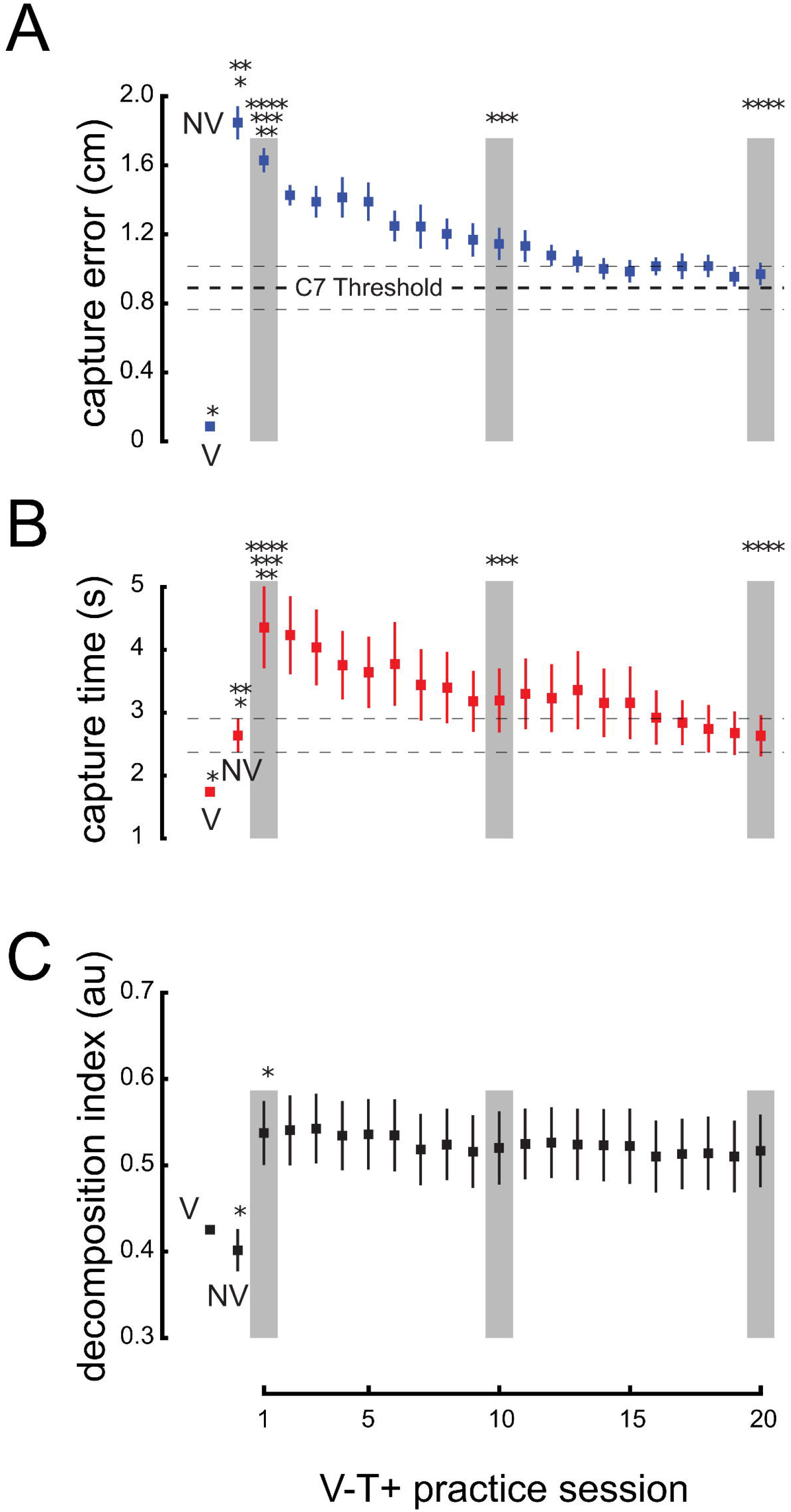
Cohort results: reach performance under single-task conditions. **Panel A)** Target capture error. The thick and thin horizontal dashed lines indicate the average pre-practice JND of VTF in dermatome C7 referenced to displacement in the workspace (0.89 ± 0.13 cm). **Panel B)** Target capture time. Horizontal dashed lines: ±1 SEM of the average cohort performance in the no-vision baseline condition on Day 1. **Panel C)** Decomposition index. Error bars: ±1 SEM in all cases. *Black* asterisks indicate a significant difference between the paired conditions. *Gray* shaded bars indicate the sessions included in the t-tests.

**Figure 4:**
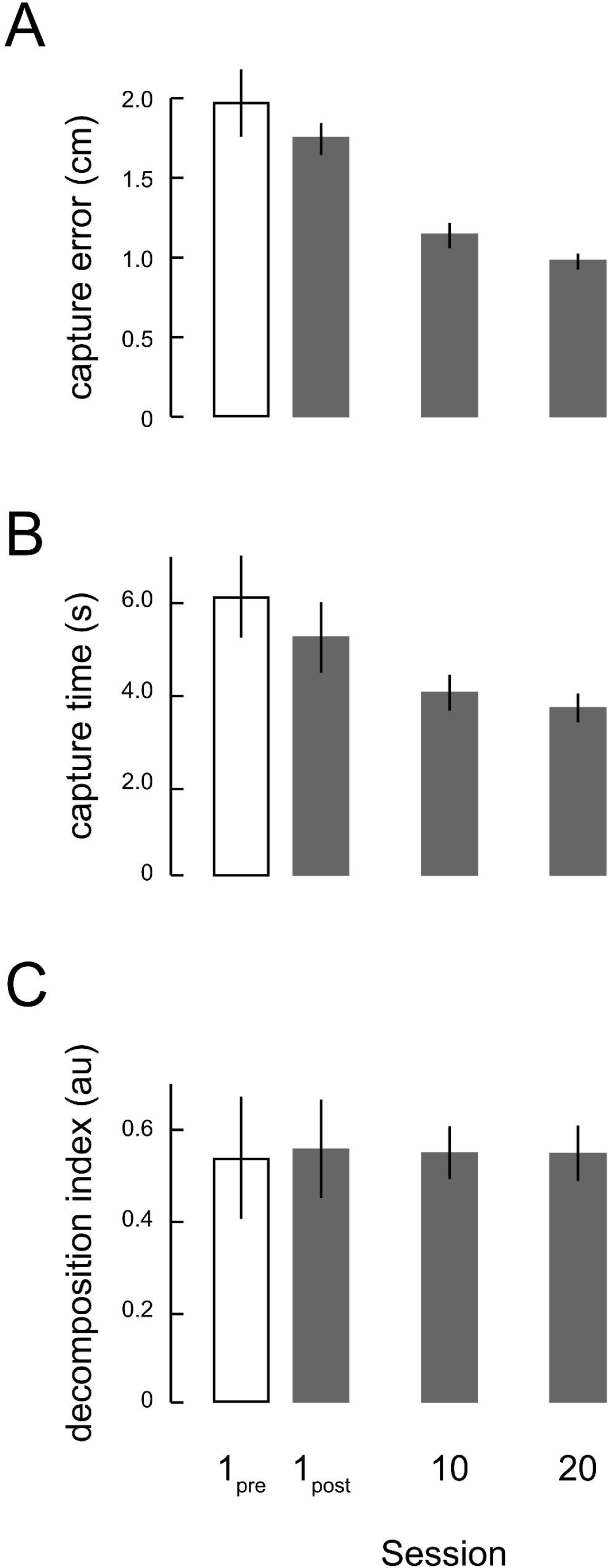
Dual-task target capture error and target capture time results for sessions 1 (pre- and post-practice), 10, and 20. **Panel A)** Group target capture error for VTF-guided reaching during dual-task blocks. **Panel B)** Group target capture time for VTF-guided reaching during dual-task blocks. **Panel C)** Group Decomposition Index for the VTF-guided reaching during dual-task blocks. Error bars indicate SEM.

Target capture errors during VTF-guided reaching asymptotically approached the limits of acuity of vibrotactile intensity discrimination (Shah et al. 2019a), as mapped onto the reachable workspace for this study (0.89 ± 0.13 cm; Fig 3A: horizontal *dashed black* line and *dashed gray* lines).

We also considered how temporal aspects of movement efficiency changed as participants practiced reaching with supplemental kinesthetic VTF. Visually guided reaches in session 1 were most efficient, requiring the least amount of time to capture the target (1.88 ± 0.26 s; V+T-; Fig 3B: “V”). Removing visual feedback (V-T-condition; Fig 3B: “NV”) resulted in somewhat longer target capture time relative to visually guided reaching (2.79 ± 1.06 s; t_14_ = 3.85, p = 0.001). Target capture time increased to 4.53 ± 2.57 seconds on initial exposure to supplemental kinesthetic vibrotactile feedback (V-T+); these capture times were significantly longer than those in the V-T-condition (t_14_ = 3.36, p = 0.002). Target capture time also decreased with practice using supplemental feedback [F_(3.06,42.8)_ = 5.08, p = 0.004], reflecting improvement in this measure of movement efficiency. Relative to initial exposure in session 1, target capture time decreased significantly by the 10^th^ practice session (3.36 ± 2.00 s; t_14_ = 2.53, p = 0.012) such that movement times were no longer statistically different from target capture time observed during V-T-condition (t_14_ = 1.10, p = 0.15). Target capture time decreased further by session 20 (2.78 ± 1.30 s; t_14_ = 3.49, p = 0.002); by the end of training, the population statistics were indistinguishable from those observed when participants reached in the V-T-condition (t_14_ = 0.013, p = 0.50).

We then considered how spatial aspects of movement efficiency may have changed with practice. Although Decomposition Index (DI) was impacted by the introduction of VTF, it did not change substantially over 10 hours of extended practice. DI observed during visually guided reaching averaged 0.42 ± 0.03 (Fig 3C: “V”). DI did not change significantly when visual feedback was removed (V-T-condition; Fig 3C: “NV”). DI = 0.39 ± 0.12; t_14_ = 1.13, p = 0.14). By contrast, DI increased markedly to 0.57 ± 0.18 during the first session of VTF-guided reaching practice. This increase relative to the V-T-condition was significant (t_14_ = 4.58, p < 0.001) reflecting the tendency of participants to move first along one cardinal axis of the vibrotactile display, then along the other. This finding was consistent with a previous report of decomposed movements during VTF-guided reaching (Risi et al. 2019). Participants persisted in using this “decomposition strategy” throughout the course of the study [F_(1.52,21.3)_ = 0.70, p = 0.47]. There was no evidence for significant reductions in DI values between the first practice session to the 10^th^ session (DI = 0.55 ± 0.21; t_14_ = 0.64, p = 0.27) or 20^th^ session (DI = 0.54 ± 0.21; t_14_ = 0.74, p = 0.24).

We also analyzed performance of the secondary choice reaction task. Participants were equally accurate in performing the target color choice test during the “Familiarization” blocks of session 1 (96.7 ± 5.0% correct), session 10 (98.3 ± 3.8% correct), and session 20 (98.9 ± 3.3% correct). Repeated measures ANOVA found no evidence for systematic difference across these sessions with regards to the accuracy of choice reaction task performance [F_(2,28)_ = 1.49, p = 0.24]. By contrast, reaction times on the secondary task decreased significantly from 753 ± 174 ms in session 1, to 647 ± 79 ms in session 10, and to 611 ± 85 ms in session 20 [F_(1.19,16.5)_ = 8.63, p = 0.007].

### Dual-task reaching and choice reaction performance

We next considered the performance of the primary and secondary tasks under dual-tasking and compared them to the performance obtained when the tasks were performed singularly. When the reaching task was performed under the dual-task condition, the target capture errors decreased because of practice on using supplemental vibrotactile kinesthetic feedback (Fig 4A), paralleling the single-task results reported above. Target capture errors averaged 1.98 ± 0.53 cm during the 1^st^ session’s pre-practice dual-task test block and averaged 1.75 ± 0.43 cm after only one session of practice. By the 10^th^ practice session, target capture errors in the post-practice block decreased to 1.15 ± 0.23 cm, and by the 20^th^ practice session they averaged 0.98 ± 0.24 cm. We performed two-way repeated measures ANOVA to examine the impact of testing condition (single-task vs. dual-task) and session number (1, 10, and 20) on primary task target capture error. We found a significant main effect of session number on target capture error [F_(1.20,16.9)_ = 56.8; p < 0.001], a main effect of testing condition [F_(1,14)_ = 6.72; p = 0.021], as well as an interaction between these factors [F_(1.36,18.8)_ = 8.24; p = 0.006] such that the initial impact of dual-tasking on target capture errors in session 1 resolved rapidly (post-practice dual-task vs. session 1 single-task reaching; t_14_ = 1.34; p = 0.10). By session 10, the difference between single-task and dual-task performance was no longer significant (t_14_=0.024; p = 0.98).

Dual-task target capture time showed a similar pattern of performance improvement (Fig 4B). Initial dual-task target capture time (session 1 pre-practice: 6.08 ± 3.57 s) remained relatively high after one session of practice (session 1 post-practice: 5.32 ± 2.68 s). By the 10^th^ practice session, average dual-task target capture time decreased to 4.12 ± 1.79 s, and by the 20^th^ session they decreased further to 3.72 ± 1.34 s. A two-way repeated measures ANOVA found significant main effects of testing condition [F_(1,14)_ = 26.5; p < 0.001] as well as session number [F_(1.39,19.4)_ = 9.28; p = 0.003] on target capture times. We observed no interaction between these factors [F_(1.13,15.8)_ = 2.27; p = 0.15]. Post-practice dual-task target capture times remained significantly higher than single-task target capture times even after 20 sessions of practice using VTF (t_14_ = 5.23, p < 0.001).

Next, we assessed how the secondary task performance impacted the tendency of participants to decompose movements onto the cardinal axes of the vibrotactile interface during VTF-guided reaching. As was observed when subjects performed the reaching task alone, participants persisted in using a “decomposition strategy” whenever they were provided supplemental kinesthetic vibrotactile feedback (Fig 4C). DI values averaged 0.542 ± 0.174 during the first session’s pre-practice dual-task test block, 0.56 ± 0.19 after one session of practice, 0.55 ± 0.20 after 10 practice sessions, and 0.55 ± 0.20 after all 20 sessions. Even though there were no changes in decomposition across the practice sessions and conditions, these results strongly suggest that subjects continued to use the supplemental kinesthetic vibrotactile feedback throughout all 20 testing sessions and did not begin to ignore it over time. Indeed, two-way repeated measures ANOVA found no significant main effects [testing condition: F_(1,14)_ = 0.04; p = 0.84; session number: F_(1.07,15)_ = 0.04; p = 0.86] or interaction effect [F_(1.22,17)_ = 2.67; p = 0.12] on DI values. Over the course of 10 hours of practice, participants did not reduce their tendency to decompose reaches onto the cardinal axes of the vibrotactile interface.

We also assessed the primary task’s impact on performance of the secondary task. Dual-tasking had minimal impact on the accuracy of secondary task performance. We performed two-way repeated measures ANOVA to examine the impact of testing condition (single-task vs. dual-task) and session number (1, 10, and 20) on secondary task accuracy. We found no change as a function of testing condition [F_(1,14)_ = 0.08; p = 0.79], no compelling evidence for change as a function of session number [F_(2,28)_ = 3.19; p = 0.06], and no interaction between these factors [F_(2,28)_ = 0.08; p = 0.93]. Participants were equally accurate in performing the target color choice test during pre-practice (97.2 ± 4.07% correct), after one session of practice (97.2 ± 4.07%), after 10 sessions (98.3 ± 2.64%), and after 20 sessions (98.9 ± 1.91%). By contrast, the impact of dual-tasking on the secondary task’s reaction time was high. A two-way repeated measures ANOVA found a significant impact of testing condition on secondary task reaction time [F_(1,14)_ = 7.27; p = 0.017], but no main effect of session number [F_(2,28)_ = 2.47; p = 0.10], and no interaction between the two factors [F_(1.41,19.7)_ = 0.07; p = 0.88]. During the pre-practice dual-task test block (session 1), choice reaction time averaged 1843 ± 956 ms. After one session of practice, choice reaction time averaged 1275 ± 709 ms (post-practice, session 1). In session 10, dual-task choice reaction time averaged 1142 ± 828 ms, while in session 20 it averaged 1154 ± 888 ms.

In a final analysis, we assessed the extent to which the primary and secondary tasks were performed serially or in parallel during the dual-task trials. Participants could use one of three strategies to coordinate the primary reaching task and the secondary choice reaction task based on whether button pressing occurred prior to, during, or after reaching to the target. Before practice on VTF-guided reaching, button presses occurred prior to reaching on 46.7% of dual-task trials, during the reaching movement on 33.3% of trials, and after reaching on 20% of trials. After 20 sessions of practice on VTF-guided reaching, button presses occurred prior to reaching on 28.9% of trials, during the reaching movement on 49.4% of trials, and after reaching on 21.7% of trials.

Across the study population, the percentage of dual-task trials wherein reaching and button pressing occurred concurrently increased significantly over the course of 20 sessions of practice (from 33.3% ± 34.1% in session 1 to 49.4% ± 38.2% in session 20; one-sided paired t-test: t_14_ = 1.90, p = 0.04). This result is consistent with the conclusion that over the course of 20 sessions of practice, VTF-guided reaching becomes increasingly resistant to dual-task interference.

## Discussion

The long-term goal of our work is to establish non-visual closed-loop control over the movements of one limb by applying supplemental vibrotactile feedback (VTF) about those movements to some other body part. Potential applications include mitigation of somatosensory impairments after stroke and enhancement of movement precision in the manual control of surgical robotics. Prior studies have shown that providing supplemental kinesthetic feedback during reaching and stabilizing movements of the arm can yield measurable improvements in accuracy and efficiency of those actions after less than 1 hour of practice in healthy individuals and in survivors of stroke (Tzorakoleftherakis et al. 2015; Krueger et al. 2017; Shah et al. 2018; Risi et al. 2019; Ballardini et al. 2021). Motivated by classical descriptions of skill acquisition, we tested the hypothesis that extended practice on VTF-guided reaching would yield performance improvements that accrue in a manner increasingly resistant to dual-task interference (Fitts and Posner 1967). Our results demonstrate that healthy participants can learn to use limb-state VTF to improve the performance of reaching in the absence of concurrent visual feedback. Target capture errors decreased across practice sessions such that performance improvements were retained from one session to the next. With extended practice, performance on VTF-guided reaching became far more accurate than reaches performed without concurrent visual and vibrotactile feedback; VTF-guided reach accuracy approached the limits of perceptual acuity for vibrotactile intensity discrimination reported previously (Shah et al. 2019a). VTF-guided reaches also improved in temporal aspects of efficiency after 10 hours of practice, achieving similar target capture times as reaches guided solely with proprioception. However, spatial aspects of efficiency did not improve with the same amount of practice; VTF-guided reaches remained as decomposed along the principal axes of the vibrotactile interface after 10 hours of practice as they were upon initial exposure. When we asked participants to perform concurrently a secondary choice reaction task, reach performance in session 1 degraded dramatically in the sense that target capture errors and target capture times both increased markedly. But whereas the impact of the secondary task on target capture errors resolved rapidly (i.e., by session 10), its impact on target capture times did not resolve by the end of testing (i.e., by session 20). The secondary choice reaction task had no impact on the spatial efficiency of reaches; decomposition index (DI) values remained high throughout all 20 sessions whether or not the secondary task was required on any given trial. We also assessed the primary task’s impact on performance of the secondary task. Whereas dual-tasking had minimal impact on the accuracy of secondary task performance, which remained consistently high from the first session to the last, the impact of dual-tasking on the secondary task’s reaction time was large; choice reaction times in the dual-task condition were effectively twice those obtained under single-task testing even though choice reaction times decreased from session 1 to session 20 in both conditions. The finding that participants tended to increase concurrency in the performance of the primary and secondary tasks is consistent with the conclusion that over the course of 10 hours of practice, the acquired skill of VTF-guided reaching increases in its resistance to dual-task interference.

### Exploiting the skin as a communications channel

The use of vibration as a mode of sensory substitution dates back at least to the work of Robert Gault, who in 1926 described a communications system that mapped audio signals from human speech into vibratory signals applied to the fingers (Gault 1926b, a). Gault showed spoken words and sentences could be correctly interpreted by hearing-impaired subjects through the sense of touch alone. After about 25 hours of practice learning to associate 58 distinct words with vibrotactile cues, subjects achieved word-for-word accuracy on interpreting 48% of novel sentences, and “correct in sense” accuracy on another 28% (Gault and Crane 1928). Gault’s findings lend plausibility to the idea of using vibrotactile cues to mitigate deficits in other sensory modalities, at least for the interpretation of symbolic information. Recent studies continue to explore how best to encode symbolic information in the distributed encoding of vibrotactile stimuli applied, for example, across the hand and across the forearm and fingers (Luzhnica and Veas 2017; Williams and Okamura 2021). Other recent studies have explored the encoding of non-symbolic information into vibrotactile stimuli for applications such as the mitigation of vestibular deficits in the control of standing balance and in the practice of desired upper extremity movements of varying degrees of complexity through vibrotactile feedback of signals related to ongoing performance (Wall et al. 2001; Lieberman and Breazeal 2007; Sienko et al. 2008, 2012; Cipriani et al. 2012; Bark et al. 2015; Krueger et al. 2017; Ballardini et al. 2020).

For a vibrotactile interface to have utility, vibratory cues must be designed and applied such that encoded information can be easily perceived and interpreted by the user [cf., Elsayed et al. (2020); Bao et al. (2018)]. Vibrotactile perception is constrained in part by the physiology of the four groups of mechanoreceptor afferents arising from cutaneous tissues and their sensitivity to vibrations of different frequencies (Johansson and Vallbo 1979). Pacinian corpuscles respond preferentially to vibratory stimuli ranging from about 40Hz to more than 400 Hz, with a frequency-dependent stimulus detection curve having a maximum sensitivity near 300 Hz (Bolanowski et al. 1988). The other mechanoreceptors are generally less sensitive to vibratory stimuli over this range of frequencies [cf., Verrillo (1992)]; these groups are collectively called “non-Pacinian” afferents and comprise rapidly adapting RA afferents, slowly adapting SA I afferents, and SA II afferents. The perceptual detection of vibrotactile stimuli depends on many factors including the site of stimulation. The fingertip is more sensitive in detecting sinusoidally vibrating stimuli than the hairy skin elsewhere on the body (Stuart et al. 2003; Bikah et al. 2008). Additional factors include: surface area of stimulation; the presence or absence of a stable surround; skin temperature; body mass index; alcohol consumption; and advancing age (Verrillo 1980; Stuart et al. 2003; Cholewiak and Collins 2003; Gandhi et al. 2011).

For practical applications of vibrotactile sensory augmentation or substitution, it is arguably more relevant to consider the perceptual discrimination of suprathreshold stimuli of different frequencies and/or intensities rather than the detection of small vibrotactile stimuli. Perceptual discrimination also depends on several factors that can impact the rate of information transfer through the skin; these include where the stimuli are presented on the body, the distance between stimulation sites (if multiple sites), the frequency of stimulation, and how task-related information is encoded into the vibrations (Mahns et al. 2006; Krueger et al. 2017; Shah et al. 2019b, a; Elsayed et al. 2020). For example, early experimental work found that the skin is able to detect a difference of about 20-30% between two constant-frequency sinusoidal stimuli [see Verrillo (1992) for a review] and that the acuity of frequency discrimination can be enhanced for stimuli that are frequency modulated in a manner emulating the time-varying stimuli used to encode speech or movement kinematics as in the current study (Rothenberg et al. 1977). Interestingly, Rothenberg and colleagues (1977) reported that the addition of an amplitude cue to the frequency cue improved discrimination performance for each person they tested, with the average difference limen (ΔF/F: i.e., JND / standard stimulus frequency) decreasing from 25% of the center frequency to 17.5%. More recently, Cipriani and colleagues (2012) reportedly attained an average vibrotactile difference limen of about 10% for stimuli presented to the forearm when the frequency and amplitude of stimuli varied coherently. Prior studies using the same stimulator technology used in the current study yielded results more in line with the results reported by Rothenberg et al. (1977), with difference limen for sequentially-presented stimuli ranging from 17% to 23% depending on which dermatome on the arm was tested (Shah et al. 2019a). These results presented in Figure 3 show why constraints on perceptual acuity may be important: when objective estimates of vibrotactile acuity are projected onto the region of the reachable workspace spanned by the map of Figure 1C, we note that the accuracy of VTF-guided reaching appears to approach asymptotically the limit of vibrotactile discrimination acuity. Future studies might beneficially explore the extent to which performance of VTF-guided reaching might be improved by efforts to improve aspects of vibrotactile perception through perceptual learning in the discrimination of vibrotactile stimuli.

### Skill acquisition in VTF-guided reaching

Motor skill is characterized by the capability to reproduce predetermined action sequences with minimal outcome uncertainty (Fitts and Posner 1967; Schmidt and Lee 2005; Magill and Anderson 2017). In their classic paper on human performance, Fitts and Posner (1967) describe three phases of learning thought to be involved in the acquisition of complex skills. These phases include an early or “cognitive” phase wherein the learner tries to understand the task in terms of its fundamental objectives, its constituent relationships between perceptual cues and available responses, and how training signals such as knowledge of results may be effectively used to improve performance. During this phase, larger levels of cognitive resources are used to determine what motor actions to produce and how to produce them. Movements are prone to performance errors and can exhibit a high degree of performance variability across repetitions. During an intermediate or “associative” phase, patterns of coordination among constituent stimulus-response units are formed, tested through practice, and re-formed when necessary. Performance errors, which are often frequent at first, are gradually eliminated and movements become more efficient. While the duration of this phase typically depends on the complexity of the task, Fitts and Posner suggest that for many tasks, ten hours of practice can suffice to reduce performance errors to acceptable levels. In the final or “autonomous” stage, component processes become less directly subject to cognitive control – and less subject to interference from other ongoing activities or environmental distractions. This last stage reflects the habituation (automatization) of the skilled action. The speed and efficiency with which the skill is performed may continue to improve in this phase, although at a continually decreasing rate. The authors note that the distinction between the three stages is rather arbitrary, and that one phase merges gradually into the next as learning progresses.

There are at least five constituent relationships to be learned when acquiring skill in using supplemental kinesthetic vibrotactile feedback to shape ongoing control of goal-directed reaches. The first is perceptual; participants must first discriminate between vibrotactile stimuli of different intensities. Discrimination of vibrotactile stimuli is subject to perceptual learning in that perceptual acuity can increase with practice when trained (Hughes et al. 1990). Second, participants must learn a mapping between how the vibrotactile stimuli vary within the vibrotactile display as the hand moves between different target locations across the workspace. Third, they must learn to invert that map to determine what pattern of VTF will correspond to a successful target capture. Fourth, participants must learn how to strategically plan and then execute desired target capture movements based on estimates of current and desired limb states derived from all available sensory sources including VTF [cf., Sober and Sabes (2003)]. Finally, subjects must use the knowledge of results provided at the end of each reach to determine how best to update each of the constituent relationships to reduce error on the next movement attempt (recall that subjects were to correct any target capture error by re-centering the cursor on the current target in anticipation of the upcoming trial). How much of the performance error on any given trial is due to errors in perception, mapping, planning or execution? How much should participants adjust each aspect of the learned skill from one movement attempt to the next? Exploration of how the brain resolves such “credit assignment” problems is an active area of ongoing research (Fu and Anderson 2008; Dam et al. 2013; McDougle et al. 2016; Rubin et al. 2021).

Although we did not test the extent to which subjects improved perceptual acuity of vibrotactile discrimination over the course of their 20 practice sessions, the asymptotic limit of target capture errors reported here aligns with the estimate of perceptual acuity obtained from a study of vibrotactile discrimination in dermatomes on the arm and hand (Fig 3A, dashed horizontal lines). We conclude therefore that perceptual learning did not contribute substantially to the pattern of results obtained in this study. It is also likely that subjects did not markedly alter how they used vibrotactile feedback to plan and execute target capture movements, given that every subject quickly (i.e., within the first few trials) adopted a “decomposition” strategy that persisted throughout all of testing. Instead, performance improvements over the 20 sessions were likely due to improvements in the ability to infer hand position from the VTF signals, and/or their ability to invert that map and infer desired VTF signals from newly presented visual targets. A future study could discern the extent to which each aspect of mapping improves over the course of practice. By occasionally asking participants to move a cursor via button presses to where they would move their hand to capture a vibrotactile target presented within the vibrotactile display, it would be possible to assess the accuracy of the subject’s VTF-to-hand-position map. Likewise, by occasionally asking participants to activate the vibrotactile display via button presses to replicate the stimuli expected for a given hand location, or for a given visual target location, it would be possible to assess the hand-to-VTF and visual-target-to-VTF maps, respectively.

We next consider reasons why participants might have decomposed VTF-guided reaches along the principal axes of the vibrotactile display. Numerous studies of sensorimotor adaptation have shown that the brain adapts reaching movements to the specific geometry of available sensory feedback (Flanagan and Rao 1995; Wei, et al. 2005; Ghez et al. 2007; Krakauer 2009; Shabbott and Sainburg 2010). For example, Flanagan and Rao (1995) examined horizontal planar reaching movements in which they manipulated the mapping between actual and visually perceived motion such that straight-line hand paths in Cartesian space resulted in curved paths in visually perceived space and *vice versa*. Participants in that study learned to make straight line paths in visually perceived space even though the paths of the hand in Cartesian space were markedly curved. By contrast, when the reaches were perceived in Cartesian space, straight line hand paths were observed. The authors concluded that reaching movements are planned in a perceptual frame of reference so as to produce straight lines in perceived space. In our study, participants made no attempt to move their hand so that the projection of its motion onto the vibrotactile display was straight and smooth even after 20 sessions of practice. This was so even though we aligned the vibrotactile interface with the Cartesian reference frame of the manipulandum to maximize correspondence between the two reference frames. It is possible that subjects persisted in decomposing their VTF-guided reaches because even after 20 sessions of practice, they had not yet formed a useful map (internal representation) of the 2D Euclidean geometry of the space spanned by the vibrotactile display. A future study should explicitly test whether an inability to form an internal representation of vibrotactile space caused decomposition by determining the extent to which a learned map acquired through practice on a subset of visual targets might generalize to targets not included in the training set. If decomposition persists despite a demonstrated ability to generalize, an alternate explanation would be needed.

One possible explanation would be a limitation in the availability of attentional resources as described by “bottleneck” models of attention allocation (Treisman and Gelade 1980; Bonnel and Haftser 1998; Driver and Spence 1998). Our finding that participants increased concurrency in the performance of the VTF-guided reaching task and the choice reaction task suggests that a limitation of attentional resources is not likely to be the main driver for the persistent decomposition behaviors observed all throughout practice. Since participants had sufficient attentional resources to increase concurrency during the dual-task testing of session 20, they should also have had access to those same resources in the single-task reaching blocks of that same session to increase the concurrency of motions along the two cardinal axes of the vibrotactile display. Thus, a bottleneck in the allocation of attention is not a likely explanation of our results.

Another possibility is that decomposition arises due to the unconscious attempt to mitigate a perceptual phenomenon known as *masking*, whereby the ability to sense one stimulus is degraded by the presence of another stimulus presented simultaneously or very close in time. Masking is a fundamental characteristic of sensory perception and has been observed to apply to auditory, visual, and haptic stimuli (Marcel 1983; Leek et al. 1991; Verrillo 1992; Strait et al. 2010). If masking degrades perceptual acuity in the discrimination of vibrotactile stimuli of varying intensities, participants could obtain more accurate results in VTF-guided reaching if they first moved the hand along one axis of the display to match to the desired intensity of one vibrator, and then along the other axis to finally capture the target. The potential impact of masking might be large; the acuity of discrimination between two vibrotactile stimuli was worse when the stimuli were presented simultaneously than when they were presented sequentially in time (Shah et al. 2019a). The penalty for simultaneous stimulation was a 35% increase in the just-noticeable-difference discrimination threshold relative to the sequential stimulation condition when the stimuli were presented *across* different dermatomes in the arm and a 60% increase for simultaneous stimulation at two sites *within* the same dermatome. Such results suggest that participants might have compelling motivation to decompose VTF-guided reaches along the cardinal axes of the vibrotactile display to improve accuracy in the presence of masking. To the extent that moving the hand straight to the target is desirable when using supplemental kinesthetic vibrotactile feedback to guide reaching, future studies should examine how (and to what extent) it may be possible to train users to move with spatial efficiency, as well as with accuracy and temporal efficiency.

### Limitations and future directions

A limitation of our study pertains to the design of our dual-task testing condition. Although participants were instructed to perform the primary and secondary tasks as quickly and accurately as possible, we did not otherwise encourage or pressure subjects to perform the two tasks concurrently either through explicit instruction, reward, or penalty. Consequently, the increased concurrency of the primary and secondary tasks reported here could have underestimated the potential increase in concurrency due to extended practice on the primary task. Another limitation pertains to our study’s focus solely on horizontal planar reaching movements. For supplemental kinesthetic feedback to be useful in a real-world scenario, movements within a 3D workspace must be considered. Adding a third pair of vibration motors to the interface to map a third movement dimension does not pose any real technical challenges. However, the vibratory cues provided by a 3D vibrotactile sensory interface may prove difficult for participants to decode in real-time. Relatedly, our study relied on the instrumentation embedded within the planar manipulandum to infer hand location. Practical use of future supplemental kinesthetic vibrotactile feedback systems will require development of wearable limb state estimators using low-cost technology such as inertial measurement units (van der Linden et al. 2009; Lee et al. 2011). In any event, future experimental efforts are warranted to assess the utility and usability of a 3D limb state estimator and vibrotactile sensory interface to enhance performance of goal-directed behaviors in 3D, especially those related to key activities of daily living such as eating, dressing, and grooming.

## ACKNOWLEDGEMENTS

We thank Dr. Kristy Nielson for her consultancy on the validity of the dual-task paradigm and Giulia Ballardini for development of the manipulandum used in this study.

